# Molecular basis of SPRTN activation by DNA and ubiquitin

**DOI:** 10.64898/2026.06.16.732584

**Authors:** Hyun-Seo Kang, Sophie Dürauer, Christian Renz, Yuka Machida, Helle D Ulrich, Yuichi J Machida, Michael Sattler, Julian Stingele

## Abstract

The promiscuous metalloprotease SPRTN is the key enzyme for proteolytic repair of DNA-protein crosslinks (DPCs). To prevent uncontrolled SPRTN activity, its activation must be tightly regulated. Using NMR spectroscopy and *in vitro* reconstitution, we elucidate the molecular basis of SPRTN’s activation by DNA and ubiquitin. We identify an autoinhibitory mechanism governed by intramolecular electrostatic interactions between a negatively charged linker helix and SPRTN’s positively charged DNA-binding domains. DNA relieves this autoinhibition by competitively displacing the linker from the DNA-binding domains, thereby inducing a conformational shift to an open, catalytically active state. This open state enables ubiquitin binding to SPRTN’s protease domain, further stabilizing the active conformation. Disruption of the interaction between the autoinhibitory linker and the DNA-binding domains locks SPRTN in a constitutively open state, resulting in enhanced protease activity. Collectively, our data reveal how DNA and ubiquitin cooperate to convert SPRTN from an autoinhibited conformation into its active state.

## INTRODUCTION

DNA-protein crosslinks (DPCs) are highly toxic DNA lesions formed through the covalent attachment of proteins to DNA. Owing to the wide variety of proteins involved, the underlying DNA structures, and the chemical diversity of the crosslink itself, DPCs constitute an exceptionally heterogeneous and structurally complex class of DNA damage^1,2^. DPCs are induced by endogenous sources, including reactive metabolites and malfunctioning topoisomerases, as well as exogenous agents such as ionizing radiation and chemotherapeutic agents^2,3^. Due to their bulky nature, DPCs obstruct essential nuclear processes, most notably DNA replication and transcription^4–8^, thereby posing a severe threat to genome integrity. Consequently, efficient DPC repair is essential for maintaining cellular homeostasis and genome stability.

Most DPC repair pathways require proteolytic processing of the crosslinked protein to enable downstream DNA repair^3,9,10^. In addition to proteasomal degradation, the central enzymes mediating DPC degradation are members of the conserved SPRTN/Wss1 metalloprotease family^11,12^. In mammals, SPRTN targets DPCs in both replication-coupled and replication-independent global-genome DPC repair pathways^13–19^. SPRTN-dependent DPC processing is essential for genome maintenance and cellular viability^20,21^. Loss of SPRTN causes severe genome instability characterized by micronuclei formation, chromosomal segregation defects, and anaphase bridges^20^. Hypomorphic germline mutations in *SPRTN* lead to expression of truncated variants lacking substantial portions of the *C*-terminal region (SPRTN^ΔC^), causing the rare hereditary disorder Ruijs-Aalfs syndrome (RJALS)^21^. RJALS is characterized by developmental abnormalities, early-onset hepatocellular carcinoma, and features of premature aging^21,22^. These phenotypes are at least in part driven by STING-dependent inflammatory signaling, which is triggered by cGAS activation in response to the accumulation of cytosolic DNA resulting from deficient DPC repair^23^.

To perform its essential role in DPC repair, SPRTN must process a wide range of protein adducts. The enzyme contains an *N*-terminal conserved SprT domain (aa28–214) composed of a promiscuous metalloprotease domain (PD) and a zinc-binding domain (ZBD) that recognizes single-stranded DNA (ssDNA)^24^ (Fig. 1a). On the face opposite the active site, the SprT domain additionally contains a ubiquitin-binding interface (USD, Ubiquitin-binding interface at the SprT Domain)^25,26^. The SprT domain is followed by the basic region (BR), a second DNA-binding domain specific for double-stranded DNA (dsDNA)^27^. Together, the SprT domain and BR constitute the minimal SPRTN construct that retains activity (SprT-BR, hereon named PD-ZBD-BR, aa28–245) ^25^. The largely unstructured *C*-terminal region, which is absent in RJALS-associated SPRTN variants, contains several additional protein-protein interaction motifs, including a SHP-box for interaction with p97^28,29^, a PIP-box mediating PCNA binding^30^, and a ubiquitin-binding zinc finger (UBZ)^30^ (Fig. 1a).

**Figure 1.**
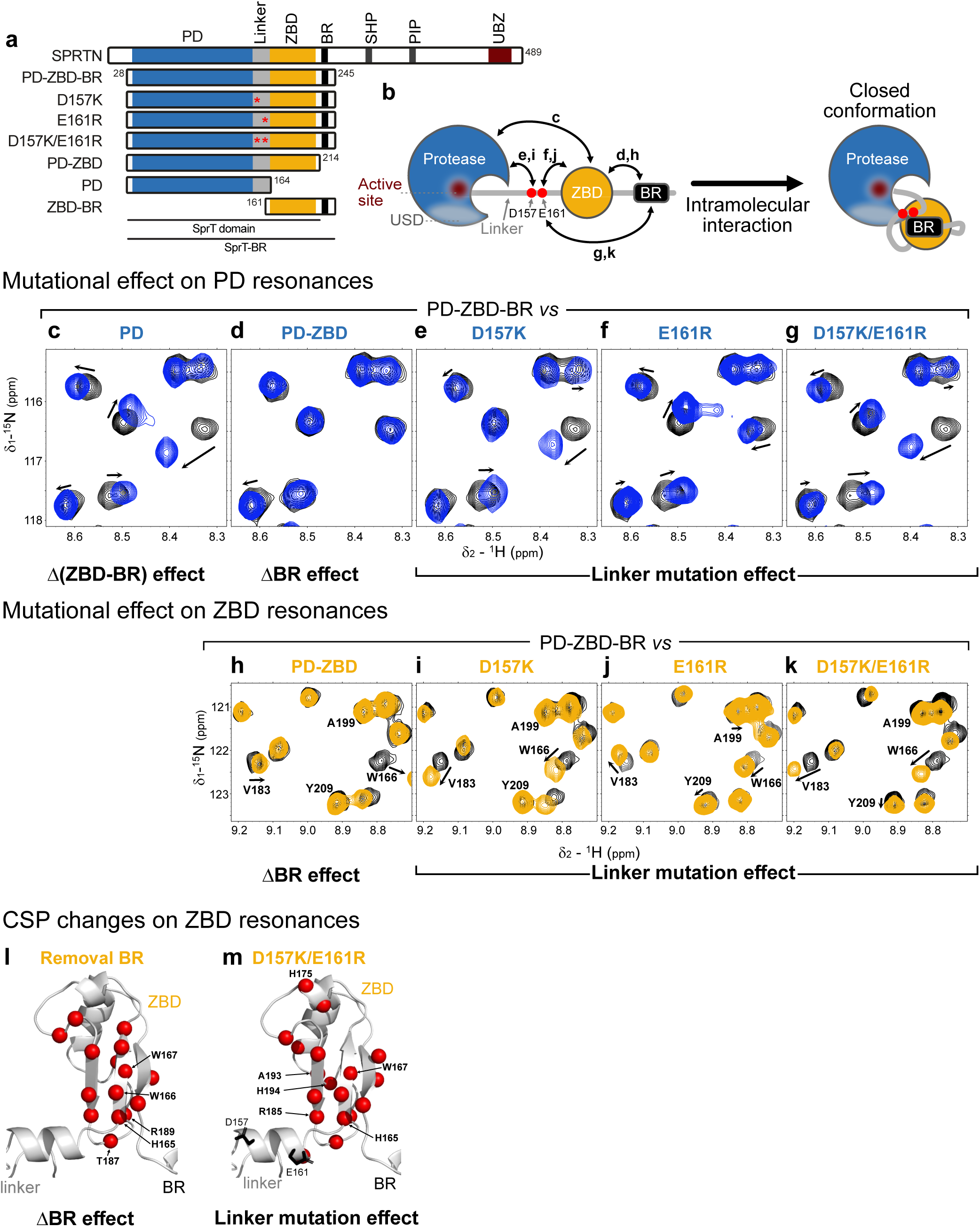
Intramolecular interactions in SPRTN. **a** Schematic of SPRTN variants, showing the minimal active construct SprT-BR (aa28-245), featuring protease domain (PD), zinc-binding domain (ZBD) and basic region (BR), named PD-ZBD-BR, and BR- or ZBD-BR-truncated variants, named PD-ZBD and PD, respectively. Single linker mutant, with amino acid substitution D157K or E161R and double linker mutant with D157K and E161R (marked by red asterisks) substitution, named accordingly. For SPRTN also the SHP box (p97 binding), the PIP box (PCNA binding) and the ubiquitin-binding zinc finger (UBZ) are indicated. **b** NMR-identified intramolecular interactions are shown as schematics with the respective spectral comparisons and domain arrangements accordingly depicted in a compact closed conformation. The active site and ubiquitin binding interface at the SprT domain (USD) are highlighted. **c-g** NMR spectra superimposition of PD-ZBD-BR (black) to various deletion and linker point mutation constructs (blue), PD (protease domain only) (**c**), PD-ZBD (**d**), D157K (**e**), E161R (**f**), D157K/E161R (**g**), for protease domain resonances to show the mutational effects on the protease domain (see Supplementary Fig. 1b-f for full spectra). Changes are shown with arrows in the spectra. **h-k** NMR spectra superimposition of PD-ZBD-BR (black) to various deletion and linker point mutations constructs (yellow), PD-ZBD (**h**), D157K (**i**), E161R (**j**), D157K/E161R (**k**), for ZBD-BR region resonances to show the mutational effects on the ZBD-BR (see Supplementary Fig. 1b-f for full spectra). Changes are shown with arrows and assignments in the spectra. **l-m** Chemical shift perturbation (CSP) of PD-ZBD-BR vs PD-ZBD (removal BR) (**l**) and D157K/E161R (**m**), larger than 0.015 ppm are shown with red spheres in the ZBD structure predicted by AlphaFold3 (see Supplementary Fig. 1g-j for full CSP analyses). Five residues with the highest CSP values are labeled. Linker mutation sites (D157K and E161R) are shown with sticks in black.

Several regulatory mechanisms that control SPRTN’s promiscuous protease activity have been identified. SPRTN is inactive in the absence of DNA but undergoes rapid activation upon engagement of its two DNA-binding domains, the ZBD and BR, with ssDNA and dsDNA, respectively^14,18,27^. Consequently, SPRTN is selectively activated by DNA structures containing both single- and double-stranded features, such as ssDNA-dsDNA junctions that arise when replicative DNA polymerases stall at DPCs^16,27^. A second layer of regulation is provided by modification of the protein adduct with ubiquitin. Although not strictly required for SPRTN activity^16^, DPC ubiquitylation strongly enhances DPC cleavage *in vitro* and in cells^13,25,26^ and ubiquitylated DPCs accumulate in the absence of SPRTN activity^31^. SPRTN is thought to be recruited to ubiquitylated DPCs through its *C*-terminal UBZ domain^13,26^, while ubiquitin binding to the USD has been proposed to enhance SPRTN’s catalytic activity by stabilizing an open active conformation of the protease^25,26^. However, the structural mechanisms that keep SPRTN inactive in the absence of DNA and trigger its activation upon DNA and ubiquitin binding remain poorly defined.

Here, we dissect mechanisms of intramolecular autoinhibition controlling the DNA-dependent conformational activation of SPRTN. Using NMR spectroscopy, we show that DNA binding induces an open SPRTN conformation. We further show that conformational transitions between open and closed conformations are controlled by electrostatic interactions between the positively charged BR and a highly acidic linker region connecting the protease domain and the ZBD. Disruption of these interactions leads to a constitutively open conformation and enhances SPRTN activity. Furthermore, we demonstrate that the DNA-induced conformational transition is required for ubiquitin binding to the USD and, thus, the subsequent stabilization of the active state. Together, our findings delineate the structural mechanism underlying DNA-mediated SPRTN activation and refine current models of SPRTN regulation.

## RESULTS

### An acidic linker mediates intramolecular domain interactions in SPRTN

Our previous NMR measurements indicated that the BR interacts intramolecularly with the ZBD in the context of a ZBD-BR construct lacking the PD^27^. In addition, molecular dynamics (MD) simulations suggested dynamic interactions between the PD and the ZBD in the absence of the BR^25^. However, whether these interactions contribute to the autoinhibited conformation of SPRTN remained unclear. To address this question, we analyzed a series of deletion and point mutant variants of PD-ZBD-BR by NMR spectroscopy (Fig. 1a-g and Supplementary Fig. 1a-f).

To assess the contribution of the BR for these intramolecular interactions, we first compared PD-ZBD-BR with PD-ZBD (aa28-214) (Fig.1a). Consistent with our previous observations^27^, deletion of the BR primarily caused spectral changes for a cluster of residues on the β-sheet of the ZBD, suggesting dynamic BR to ZBD interactions in the apo state (Fig. 1d,h and Supplementary Fig. 1b). Importantly, a small subset of PD resonances was affected upon deletion of the BR (Supplementary Fig. 1b, circles), indicating that the BR also contacts the PD. To map the effects of ZBD interactions with the PD, we generated a PD-only construct (PD, aa28-164) and compared its spectrum to that of PD-ZBD-BR (NMR spectra of PD-ZBD-BR vs PD). Removal of both DNA-binding domains induced large spectral changes for amide signals in the PD region (Fig. 1c and Supplementary Fig. 1c), indicating substantial interactions between PD and ZBD. Together with the significant spectral changes in the ZBD in an isolated ZBD-BR context compared to PD-ZBD-BR, these data provide evidence for ample intramolecular interactions between PD, ZBD, and BR in the SPRTN apo state (Fig. 1b).

Next, we sought to identify key contacts mediating these interactions. We first explored the linker region (aa151-164) between the PD and the ZBD, which is predicted to form a short acidic helix containing two Asp-Glu patches (D157/E158, D160/E161). We hypothesize that this linker might engage in intramolecular electrostatic-driven interactions with the positively charged ZBD and BR. To test this hypothesis, we introduced charge-reversal mutations within the linker, either as single linker mutation (D157K or E161R) or double linker mutation (D157K/E161R) (Fig. 1a). Spectral comparisons of D157K, E161R and D157K/E161R to PD-ZBD-BR showed significant differences for these linker variants for both PD and ZBD resonances, noticeably more for the D157K/E161R (Fig. 1e-g, for PD resonances, 1i-k, for ZBD resonances and Supplementary Fig.1d-f). Interestingly, the effects on the PD resembled those that we observed upon comparing PD with PD-ZBD-BR (Fig. 1c and Supplementary Fig. 1c). This suggests that altering the charge of the acidic linker weakens intramolecular contacts and induces conformational shifts towards the open state of the protein. Likewise, linker mutations affected residues in the β-sheet of ZBD (Fig. 1i-k--m and Supplementary Fig. 1h), reminiscent of PD- and/or BR-binding interfaces on ZBD (Fig. 1h,i and Supplementary Fig. 1g). This suggests that both ZBD to and BR interactions are disrupted by linker mutations. Of note, complete removal of the acidic linker rendered the protease domain unstable during purification, indicating that the linker constitutes an integral structural element of the protease domain.

Together, these data identify the acidic linker as a key modulator of intramolecular interactions between the PD, ZBD, and BR, supporting a model in which linker-mediated electrostatic contacts stabilize a compact closed conformation of SPRTN in the apo state.

### Cooperative intramolecular interactions stabilize the closed SPRTN conformation

Next, we investigated the correlation between different domain deletions and point mutations. Here, PD-ZBD-BR represents the compact closed conformation, retaining all the observed intramolecular interactions described above, while the PD-only construct represents the PD in the open conformation. When overlaying the NMR spectra of PD-ZBD-BR, PD-ZBD, PD, and the linker mutations (D157K, E161R and D157K/E161R), most signals superimpose well (Supplementary Fig. 2), indicating that the overall structure is not affected by the introduced point mutations or domain deletions. However, a subset of resonances showed consistent chemical shift changes in the mutant variants. Interestingly, many of these resonances are gradually shifted following a linear trend from PD-ZBD-BR (closed) to PD (open) (Fig. 2a-b and Supplementary Fig. 2). This indicates additive contributions of the intramolecular interactions for shifting populations between two states, i.e. fully closed and fully open conformations. NMR signals of the PD show the largest chemical shift differences when compared to PD-ZBD-BR (Fig. 1c and Supplementary Fig.1c), while removing the BR yields only minor chemical shift changes in the PD (Fig. 2a and Supplementary Fig. 2). However, chemical shifts changes significantly towards the PD only (open state) when the linker mutations (D157K or E161R) were introduced (Fig. 2a-b). This trend is even more pronounced for the tandem linker mutation (D157K/E161R), which almost completely shifts towards the open conformation (Fig. 2a-b). A few signals in the D157K/E161R mutant are shifted even further than in the PD protein (Supplementary Fig. 2), possibly due to a charge repulsion effect of the mutation. Consistently, comparable linear correlations were observed for ZBD resonances upon introduction of linker mutations (Fig. 2c-d). To identify the open-like conformation of the ZBD region, we compared the PD-ZBD-BR (closed) and variant spectra to the ZBD-BR only spectrum (open), which we refer to as ZBD-BR, lacking the PD and the acidic linker. When comparing the effect of single and double linker mutations we observe that in the single linker mutations (D157K, E161R) the amide signal of K184 in the ZBD is shifted towards the open conformation of ZBD-BR, with an even greater shift for the double mutant (D157K/E161R) (Fig. 2c-d and Supplementary Fig. 2).

**Figure 2.**
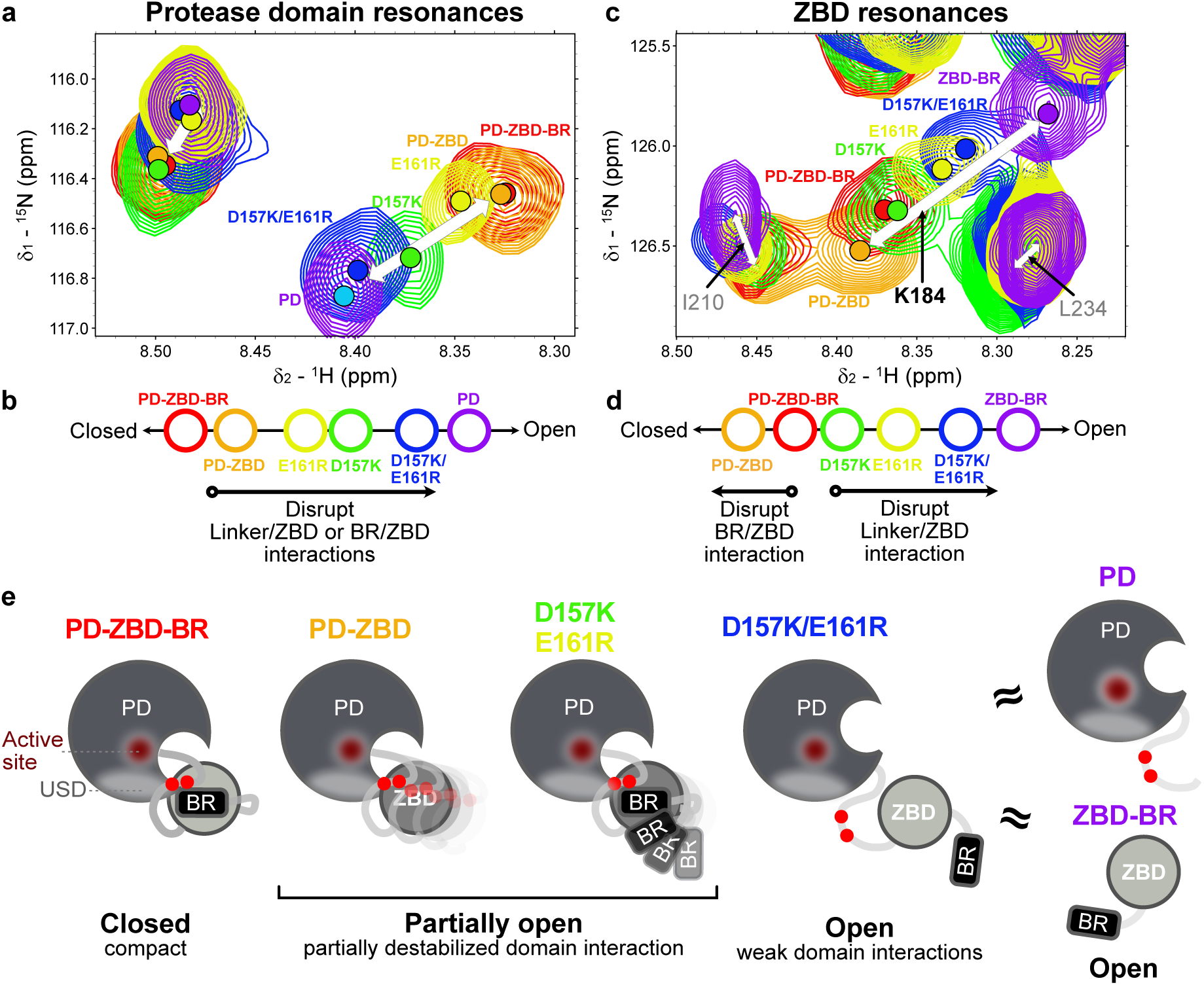
Intramolecular interaction disruptions linearly correlate between open and closed states. a-d. Superimpositions of NMR spectra (PD-ZBD-BR in red, PD-ZBD in orange, D157K in green, E161R in yellow and D157K/E161R in blue, PD or ZBD-BR in purple) for two representative regions showing PD (**a**) and ZBD (**c**) resonances for full spectra see Supplementary Fig. 2). ZBD-BR resonances are shown with assignments (K184 highlighted in bold). White arrows denote spectral changes between open and closed states. Overview of peak shifts upon disruption of intramolecular interactions are shown as schematics below for PD resonances (**b**) and ZBD resonances (**d**). **e** Schematics of the domain arrangements upon disrupting intramolecular interactions. Red spheres indicate acidic linker residues D157 and E161. The active site and ubiquitin binding interface at the SprT domain (USD) are highlighted.

The overall patterns of NMR spectral change observed in both PD and ZBD regions clearly indicate a gradual shift from closed to open conformations as intramolecular interactions are disrupted by point mutations in the linker region. PD-ZBD and single linker mutations (D157K or E161R) partially release ZBD and ZBD-BR from the PD, while the double mutant (D157K/E161R) induces an even more pronounced conformational shift towards the fully open state (Fig. 2e).

These results identify SPRTN’s acidic linker as a key structural element that enforces SPRTN’s closed apo state by pulling the positive charged DNA-binding regions of ZBD and BR close to the PD (Fig. 2e). Given the close proximity of intramolecular interfaces to the catalytic active site, we speculate that these dynamic interactions may sterically hinder access to the catalytic site and thereby dampen SPRTN’s proteolytic activity.

### Intramolecular interactions autoinhibit SPRTN proteolytic activity

Having established that linker mutations shift SPRTN toward open conformations, we next asked whether this correspondingly affects SPRTN’s proteolytic activity. To this end, we generated a defined DPC substrate by incubating the catalytic SRAP domain of HMCES (HMCES^SRAP^) with a fluorescently-labeled ssDNA-dsDNA oligonucleotide containing an abasic (AP) site^17,32^. HMCES forms covalent crosslinks with AP-sites in cells to prevent their breakage and therefore represents a relevant SPRTN substrate^33^. HMCES^SRAP^-DPCs were incubated with PD-ZBD-BR variants in presence of the helicase FANCJ, which partially unfolds the DPCs to enable SPRTN-dependent DPC proteolysis^17^.

Indeed, all linker variants displayed increased cleavage of unmodified DPCs compared with the wild type (WT) PD-ZBD-BR enzyme (Fig. 3a, compare lanes 3-5 (WT) with lanes 6-8 (D157K), lanes 9-11 (E161R) and lanes 12-14 (D157K/E161R). Similar effects were observed upon introduction of linker mutations into full-length SPRTN (Supplementary Fig. 3a). In agreement with the shift towards open conformations observed by NMR spectroscopy, these findings demonstrate that the disruption of linker-mediated intramolecular interactions enhances SPRTN’s proteolytic activity, likely by releasing the steric hinderance.

**Figure 3.**
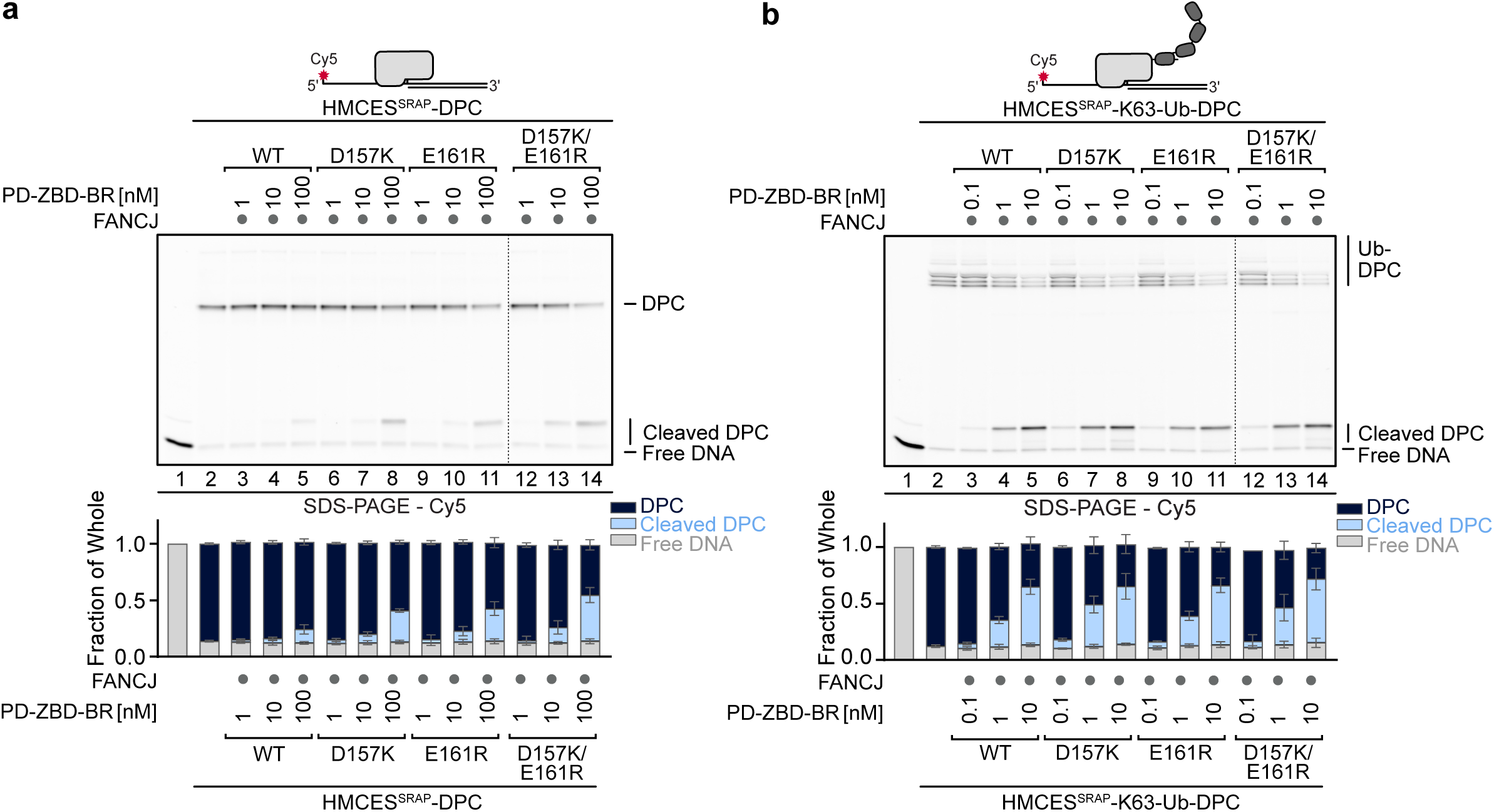
Intramolecular interactions autoinhibit SPRTN proteolytic activity a-b. HMCES^SRAP^-DPCs (**a**) or HMCES^SRAP^-K63-Ub-DPCs modified with approximately 5-8 Ub moieties (**b**) (10 nM) were incubated alone or in the presence of FANCJ (100 nM) and indicated concentrations (0.1-100 nM) of PD-ZBD-BR variants for 1 h at 30°C. Quantification: bar charts represent the mean ± SD of four (**a**) or three (**b**) independent experiments.

Because DPC ubiquitylation substantially stimulates cleavage by SPRTN *in vitro*^25,26^, we next asked whether constitutive opening of SPRTN further enhances cleavage of ubiquitylated substrates. We therefore generated K63-ubiquitylated HMCES^SRAP^-DPCs as previously described^25,34^, and assessed their cleavage by different SPRTN variants. In contrast to unmodified DPCs, all linker mutants cleaved ubiquitylated DPCs with efficiencies comparable to wild type SPRTN, in both PD-ZBD-BR (Fig. 3b) and full-length constructs (Supplementary Fig. 3b), indicating that DPC ubiquitylation can overrule the autoinhibited state of the PD-ZBD-BR module.

Consistent with their ability to process DPCs *in vitro,* SPRTN linker variants were able to rescue the loss of endogenous SPRTN in cells. We complemented conditional *Sprtn^F/−^CreER^T^*^2^ knock-out mouse embryonic fibroblasts (MEFs) with different human SPRTN-Strep variants (WT, D157K, D157K/E161R) or empty vector (EV) (Supplementary Fig. 3c-e). Loss of endogenous mouse *Sprtn* upon treatment with 4-hydroxytamoxifen (4-OHT) induced a pronounced growth defect, whereas expression of human SPRTN linker variants (Supplementary Fig. 3c) efficiently restored viability.

Together, these results demonstrate that disruption of linker-mediated autoinhibition enhances SPRTN activity toward unmodified DPCs. However, this effect is lost on ubiquitylated substrates, suggesting that DPC ubiquitylation exerts such a strong activating effect by itself, that constitutive opening of the enzyme provides no additional stimulation.

### Conformational states of SPRTN correlate with ubiquitin accessibility

To better understand the molecular basis of the effect of linker mutations and DPC ubiquitylation on SPRTN activation, we next explored conformational states of SPRTN and their link to ubiquitin binding. Our previous work had identified the USD ubiquitin binding interface within SPRTN’s protease domain located adjacent to the acidic linker^25^. MD simulations indicated that ubiquitin binding at the USD stabilizes an open SPRTN conformation^25^. We hypothesized that the intramolecular interactions involving ZBD and BR could sterically hinder ubiquitin binding to the USD.

To test this idea, we monitored the effects of ubiquitin binding on different SPRTN conformational states by NMR spectroscopy (Fig. 4a-l and Supplementary Fig. 4a-k). Addition of a five-fold molar excess of ubiquitin to PD-ZBD-BR induced only minor spectral changes, predominantly for the PD signals in the PD-ZBD-BR spectrum (Fig. 4a and Supplementary Fig. 4a) and to a lesser extent for the ZBD-BR signals (Fig. 4g and Supplementary Fig. 4g). This indicates a weak ubiquitin engagement in the predominantly closed PD-ZBD-BR conformation (Fig. 4l).

**Figure 4.**
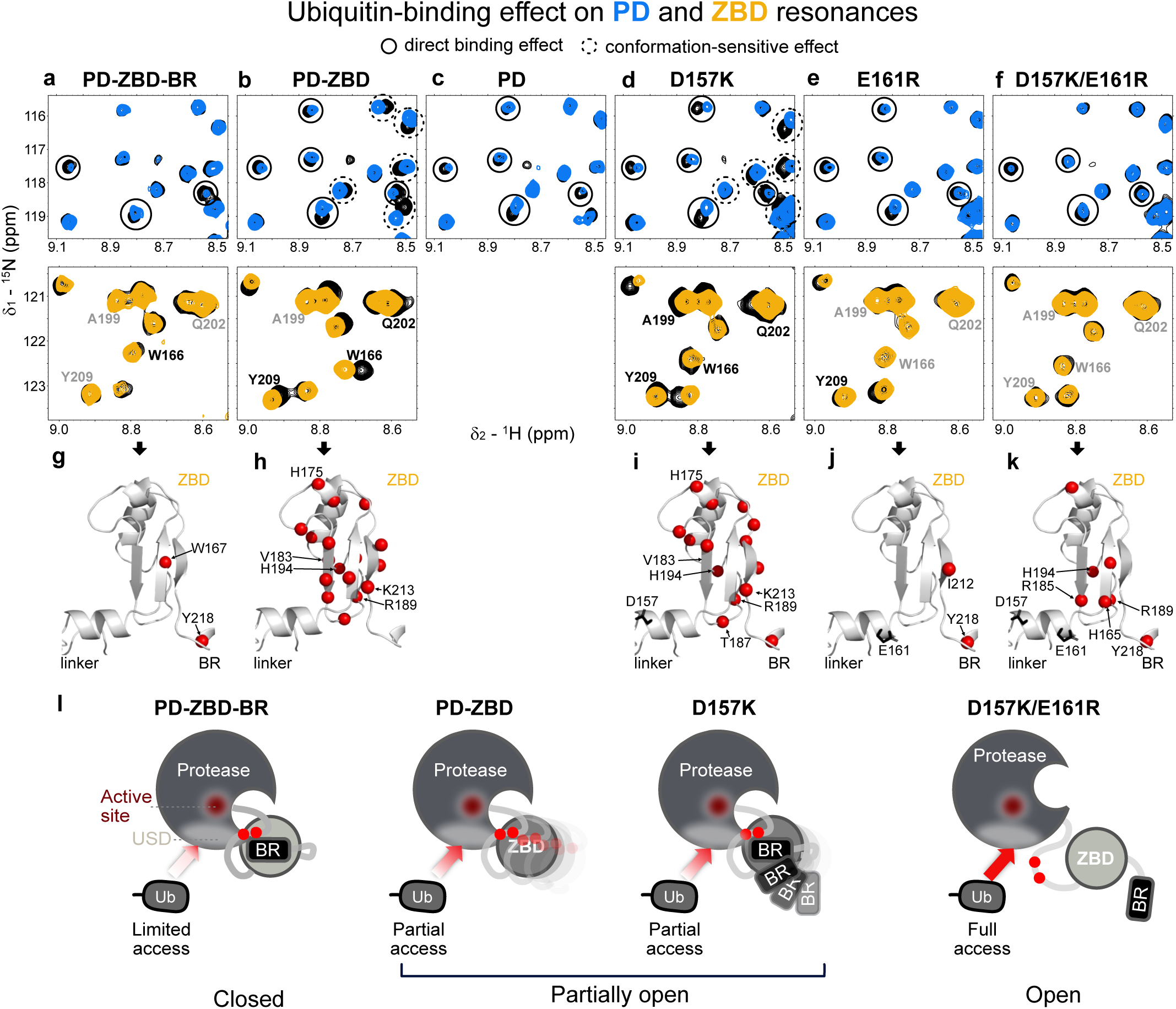
Intramolecular interactions implement an autoinhibitory effect on ubiquitin binding. a-f. NMR spectra of PD-ZBD-BR (**a**), PD-ZBD (**b**), PD (**c**), D157K (**d**), E161R (**e**) and D157K/E161R (**f**), before (black) and after (blue: protease resonances, orange: ZBD resonances) adding a 5-fold molar excess of ubiquitin (Ub). Affected resonances are shown with black circles with dashed line (conformation-sensitive change) and with solid line (direct USD-binding effect) for the PD resonances. ZBD resonances are labeled in grey (unaffected) and black (shifted or broadened) (see Supplementary Fig. 4a-f for full spectra). **g-k** For ZBD resonances, chemical shift perturbation (CSP) values larger than 0.015 ppm upon Ub addition in PD-ZBD-BR (**g**), PD-ZBD (**h**), D157K (**i**), E161R (**j**) and D157K/E161R (**k**), are shown with red spheres in the ZBD AlphaFold3 predicted structure (see Supplementary Fig. 4g-k for full CSP analyses). Top five changes are labeled with residue number.Linker mutation sites (D157K and E161R) are labeled in black. **l** Schematics of the Ub-accessibility to USD are shown upon disrupting intramolecular interactions. The active site and ubiquitin binding interface at the SprT domain (USD) are highlighted

In contrast, addition of ubiquitin to the fully open conformation (PD) caused significant shifting or broadening of specific NMR signals (Fig. 4c and Supplementary Fig. 4c), likely reflecting direct binding involving the residues in and near the USD. Surprisingly, for the partially open conformations (PD-ZBD and D157K constructs) (Fig. 4l), we observed even larger spectral changes in the presence of ubiquitin, not only for the PD but also for the ZBD-BR resonances (Fig. 4b,d,h,i and Supplementary Fig. 4b,d,h,i). A subset of the resonance changes corresponded to the direct USD binding effect, as seen similarly for the PD construct (compare Fig. 4b,d to Fig. 4c). The other subset of resonance changes, including all the changes in ZBD-BR resonances, corresponds likely to the conformation-sensitive changes, indicating transition towards more open states. The partially destabilized closed state allows ubiquitin to bind to the USD and induces conformational shifts to a more open state. This is consistent with the role of ubiquitin binding for stabilizing the open conformation as we reported previously^25^.

Lastly, we observed noticeably less spectral changes for the linker variant, E161R and D157K/E161R, unlike PD-ZBD and D157K. While we still observed some specific spectral changes for PD signals with these linker variants (Fig. 4e-f and Supplementary Fig. 4e-f), consistent with direct binding to the USD, almost no changes are observed for ZBD residues (Fig. 4j,k and Supplementary Fig. 4j,k). This is not surprising as these constructs already adopt a more open conformational state, and thus no or less conformation-sensitive resonance shifts could be observed. However, despite the consistent direct binding effect, the changes were less pronounced for NMR signals of PD (Fig. 4f and Supplementary Fig. 4f) and ZBD-BR (Fig. 4k and Supplementary Fig. 4k) compared to ubiquitin-binding effects observed in the PD-ZBD and D157K spectra (Fig. 4b,d,h,i and Supplementary Fig. 4h-i) This suggests that E161 is involved in direct interactions with ubiquitin.

In summary, the intramolecular interactions enforced by the acidic linker establish an autoinhibitory effect on ubiquitin binding (Fig. 4l). Partial disruptions of these intramolecular interactions (PD-ZBD, D157K) allow ubiquitin binding to the USD in the protease domain, which subsequently stabilizes the open conformation seen by direct binding effects and conformation-sensitive resonance changes.

### DNA binding shifts the conformational equilibrium to a more open state

Next, we asked whether DNA binding by ZBD and BR modulate the intramolecular autoinhibition within SPRTN to induce an open conformation enabling ubiquitin binding at the USD. To address this question, we monitored a set of conformation-sensitive NMR signals within PD and ZBD regions for different SPRTN variants in the presence of ubiquitin or DNA. As established above, the resonances in PD-ZBD-BR spectrum represent the closed conformation, while the resonances in PD or ZBD-BR spectra represent the open conformation for PD and ZBD-BR regions, respectively. Two distinct chemical shifts are seen for the PD resonances corresponding to the closed and open conformations represented by the resonances of PD-ZBD-BR and PD, respectively (Fig. 5a, overview in Fig. 5c). Notably, the NMR signals of the PD indicate that the compact closed state of PD-ZBD-BR is only shifted towards the open state in the presence of DNA, but not with ubiquitin. This is expected as the USD is not accessible in the apo state but only becomes available in the presence of DNA, thereby inducing a conformational shift to the open state of SPRTN.

**Figure 5.**
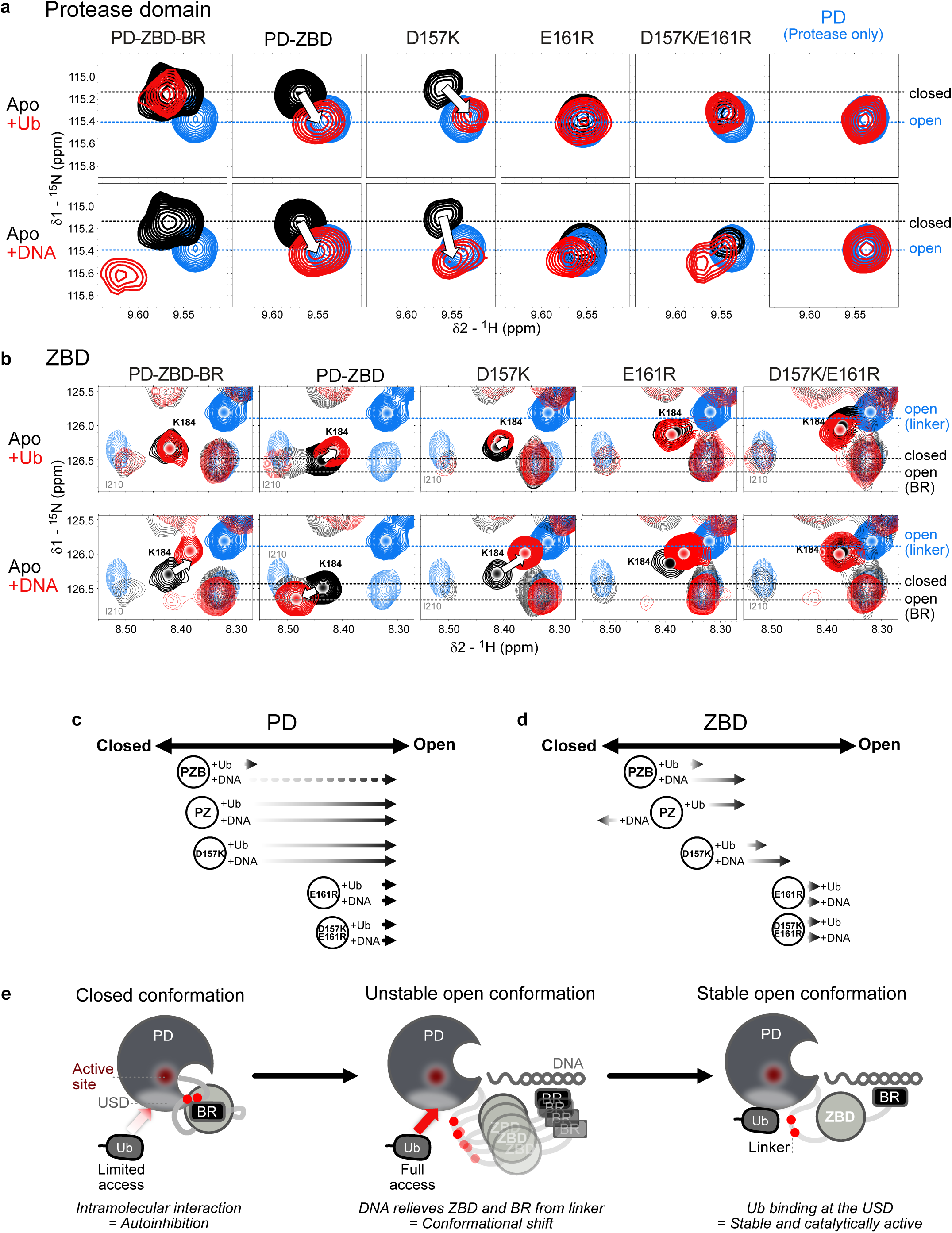
Binding to DNA or ubiquitin shifts SPRTN conformations from closed to open states. a-b. Conformation-dependent resonances in the protease domain (**a**) and ZBD (**b**) of PD-ZBD-BR (closed conformation), PD-ZBD, D157K, E161R, D157K/E161R, and PD (open conformation, for protease domain) comparing apo (black) and in the presence of a 5-fold molar excess of ubiquitin (Ub) or a 2-fold molar excess of DNA (red). Open conformation resonances (PD for protease domain, ZBD-BR for ZBD) are shown for all in cyan. For ZBD, the BR-detached open state is also indicated. **c-d** Schematics of conformational shifts from closed to open upon binding to Ub or DNA observed in the protease domain (**c**) and ZBD (**d**). **e** Model for SPRTN autoinhibition. Intramolecular interactions between the acidic linker and the DNA-binding domains, zinc-binding domain (ZBD) and basic region (BR), stabilize the closed conformation, restricting access to the active site in protease domain (PD) and limiting access to the ubiquitin binding interface at the SprT domain (USD). DNA outcompetes these interactions, and binding of ZBD and BR to ssDNA and dsDNA, respectively, triggers a conformational shift toward the open conformation, allowing full access to the USD. Ubiquitin (Ub) binding to the USD further stabilizes this open and catalytically state.

We confirmed the DNA-dependent ubiquitin binding at the USD interface observed in NMR measurements in pull-down experiments. For these experiments, we used recombinant Flag-tagged SPRTN^E112Q^ variants. We specifically chose the inactive E112Q variants to prevent activation of SPRTN during the experiment. These SPRTN variants were then incubated with K63-linked tetra-ubiquitin (K63-Ub^4^) both with and without DNA. In the absence of DNA, full-length SPRTN – which in addition to the USD contains the *C*-terminal ubiquitin-binding UBZ domain – bound efficiently to K63-Ub^4^, while SPRTN constructs lacking the UBZ domain showed strongly reduced (ΔUBZ) or no (PD-ZBD-BR) binding to ubiquitin (Supplementary Fig. 5a). Notably however, in the presence of DNA, all SPRTN variants interacted stronger with ubiquitin, including the PD-ZBD-BR. This effect was mediated by SPRTN’s USD interface, as mutation within the USD (L99S) blocked DNA-induced ubiquitin binding of the PD-ZBD-BR constructs (Supplementary Fig. 5b).

In a partially open state (PD-ZBD and D157K), both DNA and ubiquitin binding induced conformational changes to the open state. D157K/E161R resonances already superimposed well with the fully open state resonance of PD spectrum, even in the absence of DNA and ubiquitin, and thereby were not further affected by DNA or ubiquitin binding. In these comparisons, E161R behaved comparable to D157K/E161R (Fig. 5a), matching our result above that E161R is already in a more open conformation, however, with reduced ubiquitin binding affinity at the USD. In agreement, PD-ZBD-BR variants containing the E161R mutation showed reduced DNA-induced ubiquitin binding in pull-down experiments (Supplementary Fig. 5c).

The conformation sensitive residue K184 in the ZBD shows distinct chemical shifts which correlate well with conformational transition towards the open state in the presence of ubiquitin and/or DNA (Fig. 5b, overview in Fig. 5d). Although less evident compared to the PD resonances in Fig. 5a, for apo D157K/E161R, the NMR signal was clearly shifted towards the open state and unaffected by the addition of ubiquitin or DNA. Interestingly, DNA binding to PD-ZBD shifted the resonance further beyond the closed state of PD-ZBD-BR, possibly due to the absence of BR/ZBD interactions, which leads to a different chemical environment in the construct lacking the BR. These observations are confirmed with additional representative NMR signals in PD and ZBD-BR (Supplementary Fig. 6a-c).

Interestingly, the glycine resonances of G222/G224/G229 in the BR are completely broadened in the fully and partially closed conformations (PD-ZBD-BR, D157K) and are only visible in the open conformation (E161R and D157K/E161R) (Supplementary Fig. 6c). Upon addition of ubiquitin, these resonances became also visible in the D157K spectrum. This is likely due to the dynamic interaction of BR to ZBD and PD occurring in an intermediate exchange regime (micro- to millisecond dynamics) at the NMR chemical shift time scale. Thus, the absence of the BR or disrupting the BR interactions reduces the exchange broadening and renders signals observable. Consistently, in the presence of DNA (open conformation), all three glycine resonances are observed for all the constructs. This reflects, yet again, conformational dynamics of BR relative to PD and ZBD, where dynamic interaction of BR to PD and ZBD likely broaden the resonances in the closed conformation, while the BR resonances reappear upon detachment in the open conformation. We reason that the changes in the PD are generally clearer as the PD chemical shifts mainly report on the interactions of the PD with ZBD-BR, while the changes in ZBD are more complex as ZBD interactions involve both the PD and the BR (as shown in Fig. 1h).

Together, these results demonstrate that DNA and ubiquitin act on the same conformational equilibrium of SPRTN. DNA binding alone is sufficient to shift the SPRTN conformation from closed to open states, whereas ubiquitin is unable to efficiently engage with the USD in the autoinhibited conformation. Instead, DNA-induced opening exposes the USD and thereby permits ubiquitin binding, which further stabilizes the open conformation.

## DISCUSSION

Our findings indicate that the SPRTN core region (PD-ZBD-BR) involves a dynamic ensemble of autoinhibited conformations that represent a closed inactive state. This conformation can be shifted to an open state by DNA binding to the ZBD and BR, enabling efficient ubiquitin binding to the USD (Fig. 5e). These data establish a hierarchical mechanism of SPRTN activation in which DNA and ubiquitin act sequentially to relieve autoinhibition of the protease. Thereby, we provide structural mechanisms for the stimulation of SPRTN by specific DNA structures and DPC ubiquitylation^14,18,25–27^. Together with previous work, our observations allow the formulation of a model of the molecular events that occur from DPC formation to the local activation of SPRTN.

In unstressed conditions, SPRTN exists in an autoinhibited state. This closed conformational state is best considered as a dynamic ensemble of structures, consistent with line-broadening observed for NMR signals in the apo state. The autoinhibition involves electrostatic intramolecular interactions between the negatively charged linker connecting the PD and ZBD and the positively charged ZBD and BR. In the absence of DNA, both BR and ZBD interact with the PD through the linker, thereby occluding access to both the active site and the USD interface. However, once a replication fork encounters a DPC, the protein adduct undergoes ubiquitylation^8,16,35^. The ubiquitin signal is recognized by SPRTN’s UBZ, leading to the enrichment of the protease at the lesion site^13,28,29^. There, the ssDNA-dsDNA junction that arose when the replicative DNA polymerase stalled at the protein adduct^16^ outcompetes the linker for BR binding and also engages with the ZBD^27^. This conformational activation not only exposes the active site but also the USD interface (Fig. 5e). Therefore, the open conformation is further stabilized by ubiquitin binding at the USD, explaining the 100-fold activation of SPRTN by DPC ubiquitylation^25,26^.

Our study illustrates how dynamic intramolecular interactions can restrain a promiscuous protease by autoinhibition and couple its activation to multiple damage-associated signals. Such hierarchical activation mechanisms may represent a general strategy by which cells achieve high specificity while retaining the ability to rapidly respond to genome instability.

## METHODS

### Mammalian cell lines

*Sprtn^F/−^* mouse embryonic fibroblasts (MEFs) (H7) immortalized with SV40 large T and transduced with a retroviral vector expressing Cre-ER^T2^ (ref^20^) were cultured in DMEM supplemented with 10% (v/v) FBS. *Sprtn* KO was induced by treating 4×10^5^ cells with methanol (MeOH) (vehicle control) or 2 μM (Z)-4-hydroxytamoxifen (4-OHT) (Ht7904, Sigma) in MeOH for 48 h. Conversion of the floxed *Sprtn* allele (F) to the KO allele (-) was verified by PCR using WT- and KO-specific forward primers and a common reverse primer. PCR conditions were 35 cycles of 94°C for 30 s, 60°C for 30 s, and 72°C for 1 min. PCR products are 527 bp and 278 bp for the floxed and the KO alleles respectively. For exogenous expression of human SPRTN in MEFs, cells were infected with retroviral vectors produced in HEK293T/17 (CRL-11268, ATCC) by co-transfecting pMSCV.hyg-SPRTN-Strep with gag-pol and VSV-G packaging plasmids. Infected cells were selected with Hygromycin B (200 µg/mL) (10687010, Thermo Scientific) for 8 days.

Protein expression was confirmed by lysing cells in NP-40 lysis buffer (50mM Tris-HCl, pH 7.4, 150mM NaCl, 0.1% NP-40 (IGEPAL) (I8896, Sigma), 10% glycerol, 5mM EDTA (BP118-500, Fisher BioReagents), 50mM NaF, 1mM Na3VO4) supplemented with protease inhibitor cocktail (P8849, Sigma). Cell lysates (30 μg protein) were resolved on SDS-PAGE gels (4-12% Bis-Tris NuPAGE, Thermo Scientific) using MOPS buffer. Resolved proteins were subsequently immunoblotted using anti-SPRTN antibody (1:500) (6F2) (ref^39^) and anti-GAPDH antibody (1:1000) (GT239, GTX627408-01, GeneTex).

### Protein expression and purification

#### SPRTN

SPRTN variants with amino acids replacements, deletions or additions were generated using the Q5-site-directed mutagenesis kit (E0554S, Sigma). Recombinant (Flag-tagged) SPRTN (Full length, ΔUBZ, PD-ZBD-BR – WT or in combination with D157K, E161R, and E112Q amino acid replacements) protein was expressed in BL21 (DE3) *E. coli* cells and purified as previously described^25^.

Briefly, BL21 (DE3) *E. coli* cells (C600003, Thermo Scientific) were grown at 37°C until reaching OD 0.7 in Terrific broth (TB) medium (prepared with tap water). Protein expression was induced addition of 0.5 mM Isopropyl-β-D-thiogalactoside (IPTG) (I6758, Sigma) overnight at 18°C. The following day, cells were harvested, snap-frozen in liquid nitrogen and stored at −80°C. All subsequent steps were carried out at 4°C. For protein purification, cell pellets were resuspended in buffer A (50 mM HEPES/KOH pH 7.2, 500 mM KCl, 1 mM MgCl_2_, 10% Glycerol, 0.1% IGEPAL (I8896, Sigma), 0.04mg/mL Pefabloc SC (76307, Sigma), cOmplete EDTA-free protease inhibitor cocktail tablets (4693132001, Roche), 1mM Tris(2-carboxyethyl)phosphine hydrochloride (TCEP)) (HN95.3, Roth) and lysed by sonication and incubated with smDNAse (45 U/mL lysate) (MPI for Biochemistry) for 30 min prior to removal of cell debris by centrifugation at 18,000 g for 30 min. Cleared supernatant was filtered using syringe filters (PVDF, 0.22 µm) and applied to Strep-Tactin®XT 4Flow® high-capacity cartridges (2-5028-001, IBA Lifesciences). Eluted proteins were further applied to HiTrap Heparin HP affinity columns (17040701, Cytiva) before desalting using PD-10 desalting columns (17085101, Cytiva). The affinity tag was cleaved off at 4°C overnight by addition of His-tagged TEV protease with 1:10 mass ratio. Cleaved recombinant SPRTN protein was further purified by size exclusion chromatography (SEC) using a HiLoad 16/600 Superdex 200 pg column 28989335, Cytiva) equilibrated in buffer B (50 mM HEPES/KOH pH 7.2, 500 mM KCl, 10% Glycerol, 1 mM TCEP (HN95.3, Roth)). Eluted proteins were concentrated with 10 kDa cutoff Amicon Ultra centrifugal filters (UFC801096, Merck) before aliquoting, snap-freezing in liquid nitrogen and storage at −80°C.

For truncated SPRTN variants smaller than 30 kDa including PD, PD-ZBD, PD-ZBD-BR (WT, L99S, D157K, E161R, D157K+E161R) and ZBD-BR, a Strep-tagged TEV protease was used. Prior to SEC, Strep-tagged TEV protease, residual uncleaved protein and the cleaved Tag were removed by a Strep-TactinÆXT 4FlowÆ high capacity cartridges^25^.

For NMR experiments, PD, PD-ZBD, PD-ZBD-BR (WT, D157K, E161R, D157K+E161R), and ZBD-BR were expressed in ^15^N-containing media as previously described^25^. For SEC, buffer E (50 mM HEPES/KOH pH 7.2, 500 mM KCl, 1% Glycerol, 2 mM TCEP, pH 7.2) was used.

#### Mono-Ubiquitin

Recombinant mono-ubiquitin (Ub^1^) was expressed in Rosetta *E. Coli* cells (70-954-3, Sigma) and purified as previously described^25^.

#### FANCJ

Recombinant FANCJ protein was expressed in High Five cells and purified as previously described^32^.

#### HMCES^SRAP^

Recombinant HMCES^SRAP^ and HMCES^SRAP^-Ub(G76V)-3C-FKBP-His6 protein was expressed in BL21 (DE3) *E. coli* cells (C600003, Thermo Scientific) and purified as previously described^25,32^.

### HMCES^SRAP^ Ubiquitylation

Recombinant HMCES^SRAP^ was K63-ubiquitylated with a simplified Ubiquiton system^34^, based on fusions of a complete ubiquitin as previously described^25^.

### DPC cleavage assay

DPCs were generated between HMCES^SRAP^ or HMCES^SRAP^-K63-Ub^[long]^ and a 30nt Cy5-labeled forward oligonucleotide (5’-Cy5-CCCAAAAAAAAAAAdUAAAAAAAAAAAACCC-3’), as previously described^17,32^. To form ssDNA-dsDNA junctions 1 µL complementary 15nt reverse oligonucleotide (5’-GGGTTTTTTTTTTTT-3’) (12 µM in nuclease-free H_2_O) was annealed to all crosslinking reactions.

DPC cleavage by SPRTN was assessed in 10 µL reactions at 30°C for 1 h, containing SPRTN variants (as indicated – concentrations ranging from 0.1-100 nM), DPC or free DNA (10 nM) and FANCJ (100 nM). The reaction buffer comprised 17.1 mM HEPES/KOH pH 7.5, 85.6 mM KCl, 3.1% glycerol, 5 mM TCEP (HN95.3, Roth), 2.1 mM MgCl_2_, 0.12 mg/ml BSA (AM2616, Thermo Scientific) and 1 mM ATP (R0441, Thermo Scientific). Reactions were stopped with 4x LDS sample buffer (NP0007, Thermo Scientific) supplemented with 5% β-ME (M3148, Sigma) and boiling for 1 min at 95°C. Samples were resolved on SDS-PAGE gels (12% Bis-Tris BOLT, Thermo Scientific) using MOPS buffer. Gels were scanned using a BioRad Chemidoc MP system with appropriate filter settings for Cy5 fluorescence. DPC cleavage was quantified using ImageJ (v1.54f), by measuring the relative Cy5 signals of HMCES^SRAP^*-*DPCs, cleaved HMCES^SRAP^*-*DPCs and free DNA.

### Flag-pull-down assays

Reactions were performed in 50 µL on ice for 15 min, containing indicated Flag-SPRTN^E112Q^ variants (1 µM), K63-Ub^4^ (0.25 µM) (SI6304, lifesensors) and optionally dsDNA (2 µM) (orwardforward: 5’-CCTTGCTAGGACATC-3’ + reverse: 5’-GATGTCCTAGCAAGG-3’, annealed to dsDNA). The reaction buffer contained 17.7 mM HEPES/KOH pH 7.2,117 mM KCl, 3.5% glycerol, 0.5 mM TCEP (HN95.3, Roth) and 0.1% IGEPAL (I8896, Sigma). After incubation on ice, reactions were transferred to Anti-FlagÆ M2 magnetic beads (10 μL per sample) (M8823, Sigma) equilibrated with Flag-wash buffer (5 mM HEPES pH 7.2, 100 mM KCl, 2% glycerol, 0.1% IGEPAL (I8896, Sigma)). Reactions were incubated for 5 min on a rotating wheel at 4°C and washed 7 times with 500 μL Flag-wash buffer each. Bound protein was eluted with 30 µL 3xFlag Peptide (100 ng/μL) (F4799, Sigma) by incubating for 30 min on a rotating wheel at 4°C. Supernatants were collected and boiled with 4x LDS sample buffer (NP0007, Thermo Scientific) supplemented with 5% β-ME (M3148, Sigma) for 10 min at 95°C. Samples were resolved on SDS-PAGE gels (12% Bis-Tris BOLT, Thermo Scientific) using MOPS buffer and subsequently immunoblotted using anti-K63-Ub antibody (5621, cell signaling technology) or stained with a Coomassie-based protein stain (GRP1, GRP).

### Cell viability

For cell proliferation assays, 1,000 cells were seeded per well (n=8) in a 96-well plate, and cell numbers were recorded every 8 h for 3 days using Cytation 5 (BioTek) equipped with a 4x objective and the Gen5 software (ver. 3.14).

### NMR spectroscopy

All NMR samples were prepared uniformly ^15^N-labeled at concentration of 100µM in 20mM HEPES/KOH pH 7.2, 150mM KCl with 10% D_2_O, as lock signal. All NMR experiments were recorded at 308K on a 600MHz Bruker Avance NMR spectrometer, equipped with a cryogenic triple-resonance gradient probe. NMR spectra were processed using TOPSPIN (3.7) and analyzed using NMRFAM-SPARKY.

## DATA AVAILABILITY

All data supporting the findings of this study are available from the corresponding authors upon request. Source data are provided with this paper.

## CODE AVAILABILITY

No custom code or mathematical algorithms were generated or used during the current study.

## ACKNOWLEDGMENTS

We thank Dennis Beuke for help with SPRTN purifications. We are grateful to Sam Asami and Gerd Gemmecker for help with NMR measurements at the Bavarian NMR center. IMB’s Core Facility for Protein Production is acknowledged for their expert support. Research in the lab of J.S. is supported by European Research Council (ERC CoG 101124695 DECONSTRUCT) and The Vallee Foundation. H.D.U. acknowledges funding by the European Research Council (ERC AdG 101140447). We acknowledge funding by the Deutsche Forschungsgemeinschaft (J.S. and H.D.U.: Project-ID 393547839 – SFB 1361) (J.S. and M.S.: Project-ID 533767322 – EXC 3113/1, Cluster for Nucleic Acid Sciences and Technologies – NUCLEATE) (M.S.: Project-ID 325871075 – SFB1309).

## AUTHORS CONTRIBUTIONS

Conceptualization: H.S.K, S.D. M.S. and J.S. Investigation: H.S.K., S.D., Y.J.M, Y.M. and C.R. Writing – Original draft: H.S.K and S.D. Writing – Review & Editing: H.S.K, S.D., M.S. and J.S. with input from all authors. Funding Acquisition and Supervision: J.S., M.S., Y.J.M and H.D.U.

## COMPETING INTERESETS

The authors declare no competing interests.

---

**Supplementary Figure 1.**
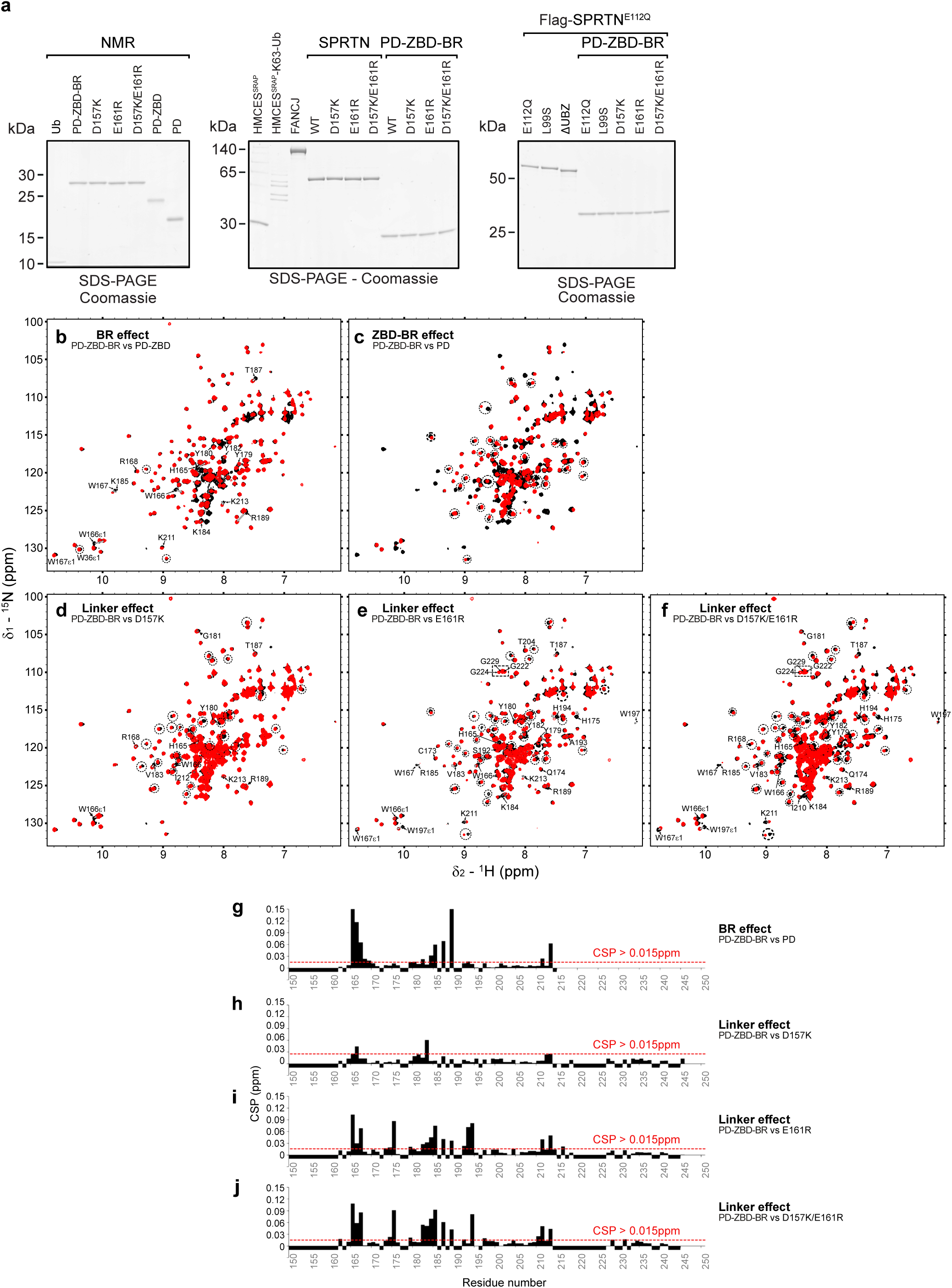
Intramolecular interactions in SPRTN. **a** Coomassie-stained SDS-PAGE gels showing equimolar amounts of purified recombinant mono-ubiquitin (Ub), PD-ZBD-BR, D157K, E161R, D157K/E161R, PD-ZBD and PD, used for NMR experiments; HMCES^SRAP^, HMCES^SRAP^-K63-Ub, FANCJ, SPRTN^WT^, SPRTN^D157K^, SPRTN^E161R^, SPRTN^D157K/E161R^, PD-ZBD-BR^WT^, PD-ZBD-BR^D157K^, PD-ZBD-BR^E161R^, PD-ZBD-BR^D157K/E161R^ used for *in vitro* DPC cleavage assays; Flag-SPRTN^E112Q^, Flag-SPRTN^E112Q+L99S^, Flag-ΔUBZ^E112Q^, Flag-PD-ZBD-BR^E112Q^, Flag-PD-ZBD-BR^E112Q+L99S^, Flag-PD-ZBD-BR^E112Q+D157K^, Flag-PD-ZBD-BR^E112Q+E161R^, Flag-PD-ZBD-BR^E112Q+D157K/E161R^ used for Flag-pull-down assays. Brightness and contrast of Coomassie images were globally adjusted. **b-f** Full NMR spectra superimposition of PD-ZBD-BR (black) to various deletion and linker point mutation constructs (red), PD-ZBD (BR removal) (**b**), PD (protease domain only) (**c**), D157K (**d**), E161R (**d**) and D157K/E161R (**e**) (related to Fig. 1c-g, respectively). Representative changes are circled in the spectra of the protease domain resonances and assignments are shown for the ZBD-BR resonances with CSP > 0.015 ppm. **g-j** CSP values are calculated for the ZBD-BR resonances, comparing PD-ZBD-BR vs PD-ZBD (**g**), D157K (**h**), E161R (**i**) and D157K/E161R (**j**), by using 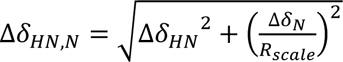, where *R*scale=6.5, as suggested (Mulder *et al.*40). Red lines denote the 0.015 ppm cutoff used for the highlighted residues in Fig. 1h-k. Negative values denote prolines or unassigned residues (related to Fig. 1l-m).

**Supplementary Figure 2.**
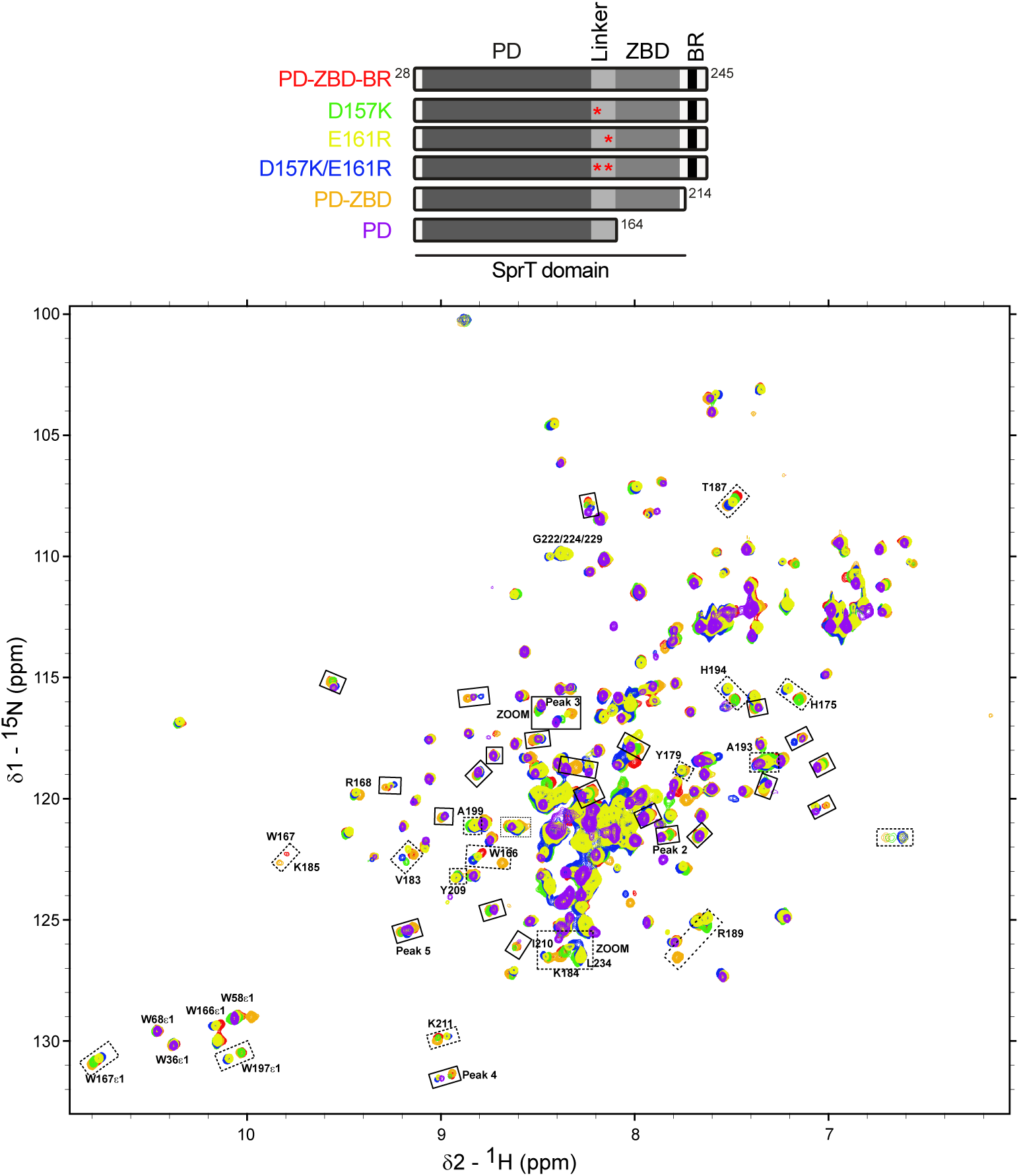
Intramolecular interaction disruptions linearly correlate between open and closed states. Schematics of SPRTN constructs (top). Superimpositions of full NMR spectra (bottom) for PD-ZBD-BR (red), PD-ZBD (orange), D157K (green), E161R (yellow) and D157K/E161R (blue), PD (purple). Boxes with solid lines denote resonance changes in the protease domain, while ZBD-BR resonance changes are shown with dashed lines and assigned. Two zoom regions shown in Fig. 2a,c are indicated. Peaks shown in Supplementary Fig. 6 are labeled.

**Supplementary Figure 3.**
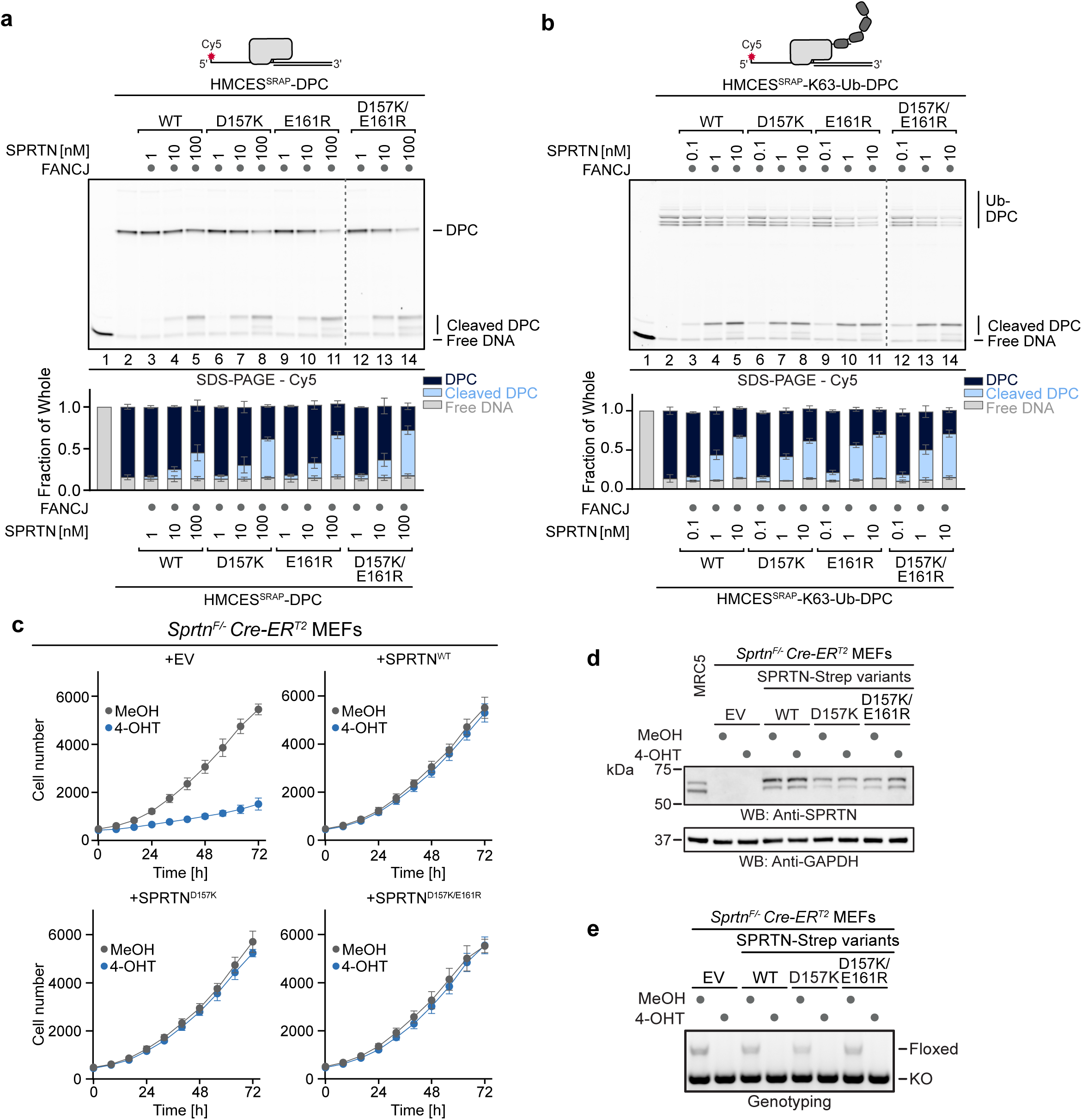
Intramolecular interactions autoinhibit SPRTN proteolytic activity. a-b. HMCES^SRAP^-DPCs (**a**) or HMCES^SRAP^-K63-Ub-DPCs modified with approximately 5-8 Ub-moieties (**b**) (10 nM) were incubated alone or in the presence of FANCJ (100 nM) and indicated concentrations (0.1-100 nM) of SPRTN variants for 1 h at 30°C. Quantification: bar charts represent the mean ± SD of three independent experiments. **c** Proliferation of *Sprtn^F/−^* Cre-Er^T2^ mouse embryonic fibroblasts (MEFs) complemented with indicated SPRTN-Strep variants or empty vector (EV, pMSCV) treated with methanol (MeOH) or (Z)-4-hydroxytamoxifen (4-OHT) (2 µM) for 48 h. After seeding, cell numbers were counted at indicated time points. Values are the mean ± SD of eight technical replicates. Shown is a representation of three independent experiments. **d** Expression of indicated SPRTN-Strep variants or EV (pMSCV) in *Sprtn^F/−^*Cre-Er^T2^ MEFs treated with MeOH or 4-OHT (2 µM) for 48 h. Whole cell lysates were analyzed by immunoblotting. **e** PCR-based genotyping of Sprtn alleles for *Sprtn^F/−^*Cre-Er^T2^ MEFs, complemented with indicated SPRTN-Strep variants or EV (pMSCV), treated with MeOH or 4-OHT (2 µM) for 48 h.

**Supplementary Figure 4.**
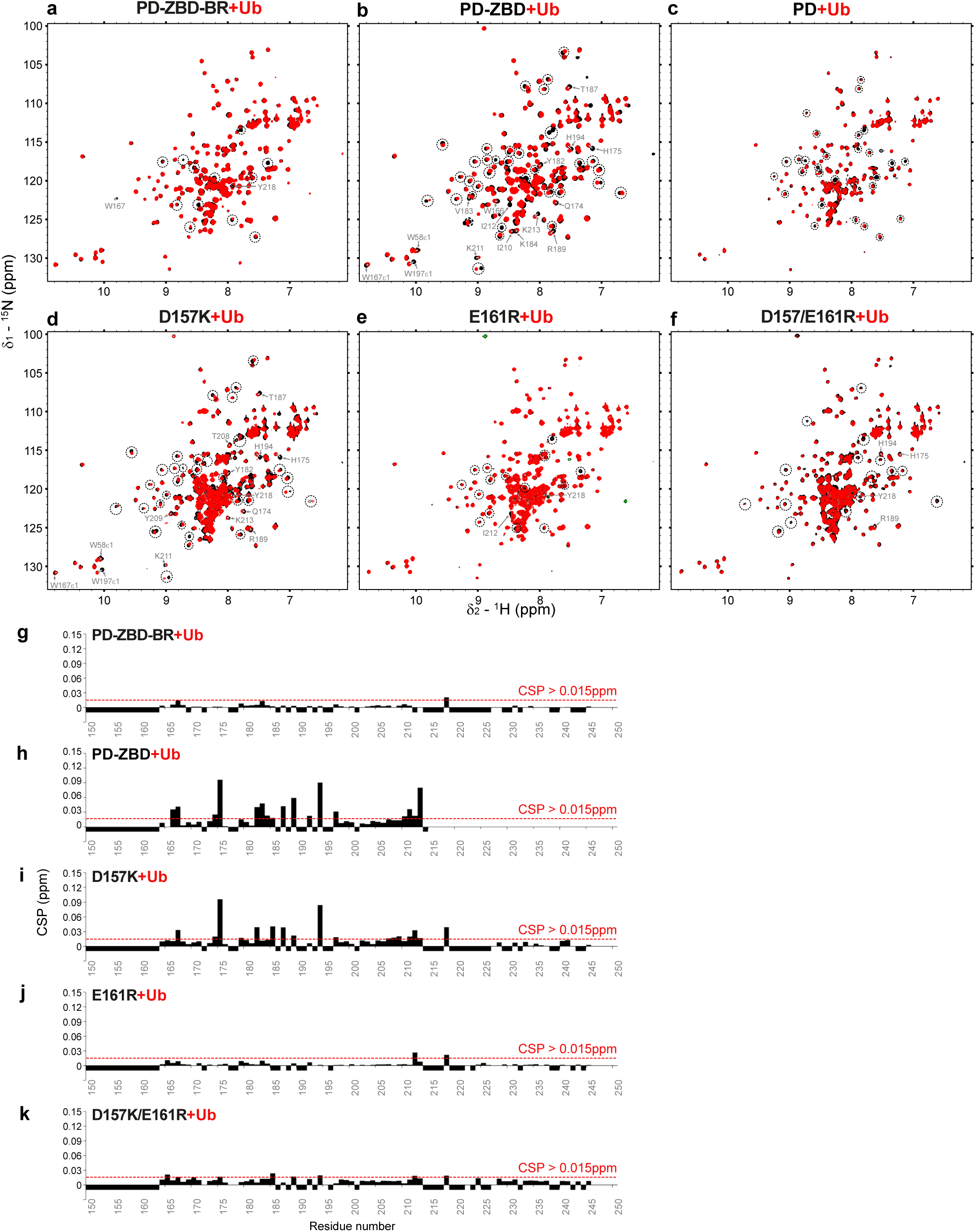
Intramolecular interactions implement an autoinhibitory effect on ubiquitin binding. a-f. Full NMR spectra of PD-ZBD-BR (**a**), PD-ZBD (**b**), PD (**c**), D157K (**d**), E161R (**e**) and D157K/E161R (**f**), before (black) and after (red) adding a 5-fold molar excess of ubiquitin (Ub). Affected resonances of the protease domain are circled. Affected ZBD resonances are labeled (related to Fig. 4a-f). **g-k** CSP analyses of the ZBD-BR resonances upon adding Ub in PD-ZBD-BR (**g**), PD-ZBD (**h**), D157K (**i**), E161R (**j**) and D157K/E161R (**k**), by using 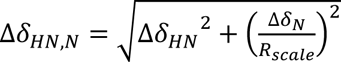, where *R*_scale_=6.5, as suggested (Mulder *et al.*^40^). Red lines denote 0.015 ppm cutoff used for the highlighted residues in Figure 4g-k. Negative values denote prolines or unassigned residues.

**Supplementary Figure 5.**
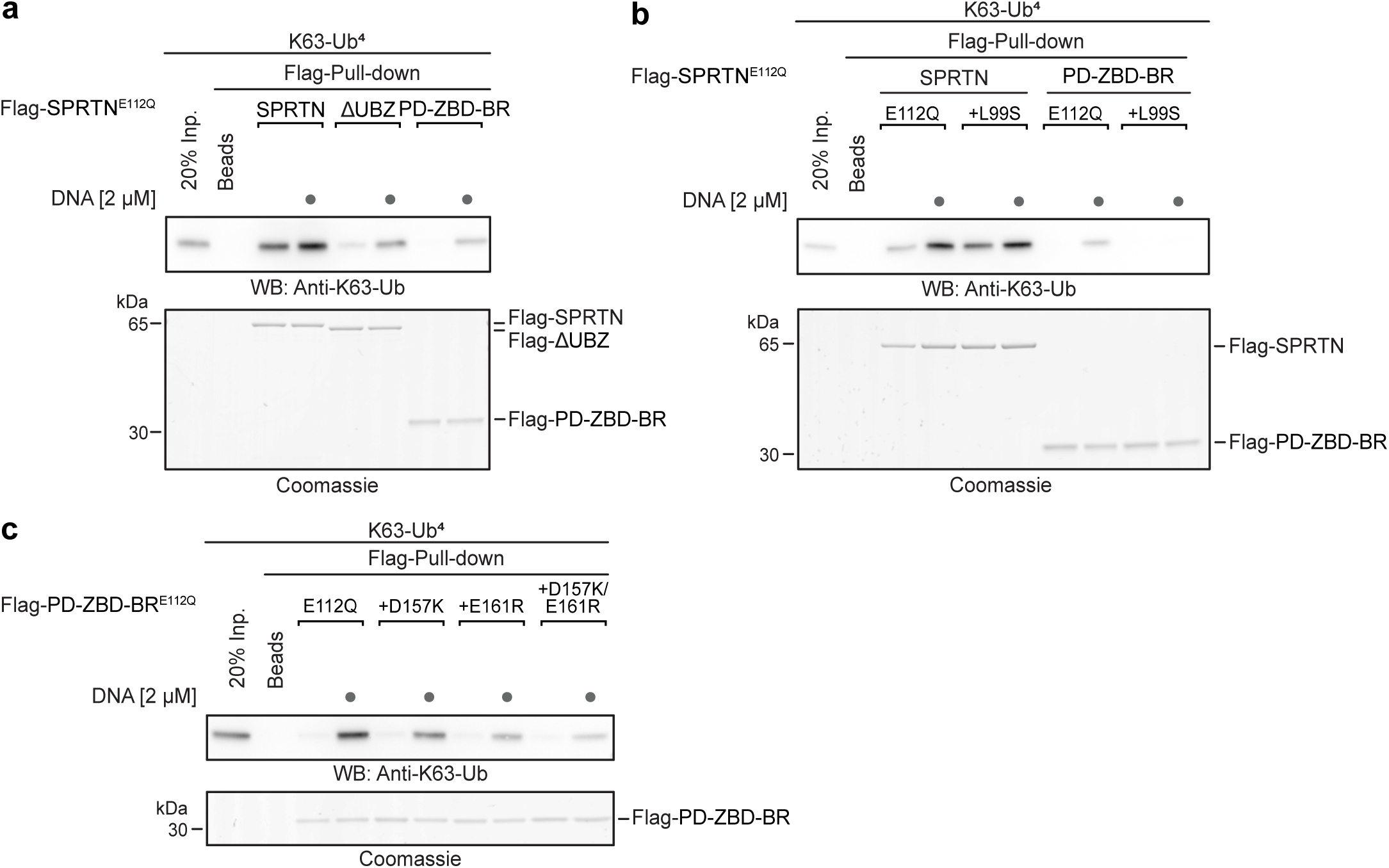
Binding to DNA or ubiquitin shifts conformations from closed to open states. a-c. K63-tetra-ubiquitin (K63-Ub^4^) (0.25 µM) was incubated alone (input sample was diluted to 20% Inp.) wor together with indicated Flag-SPRTN^E112Q^ variants (1 µM) with and without DNA (2 µM) for 15 min on ice. Flag-SPRTN^E112Q^ variants were isolated by pull-down together with bound ubiquitin using Anti-Flag beads. Reactions were analyzed by immunoblotting or Coomassie staining. Brightness and contrast of Coomassie images were globally adjusted.

**Supplementary Figure 6.**
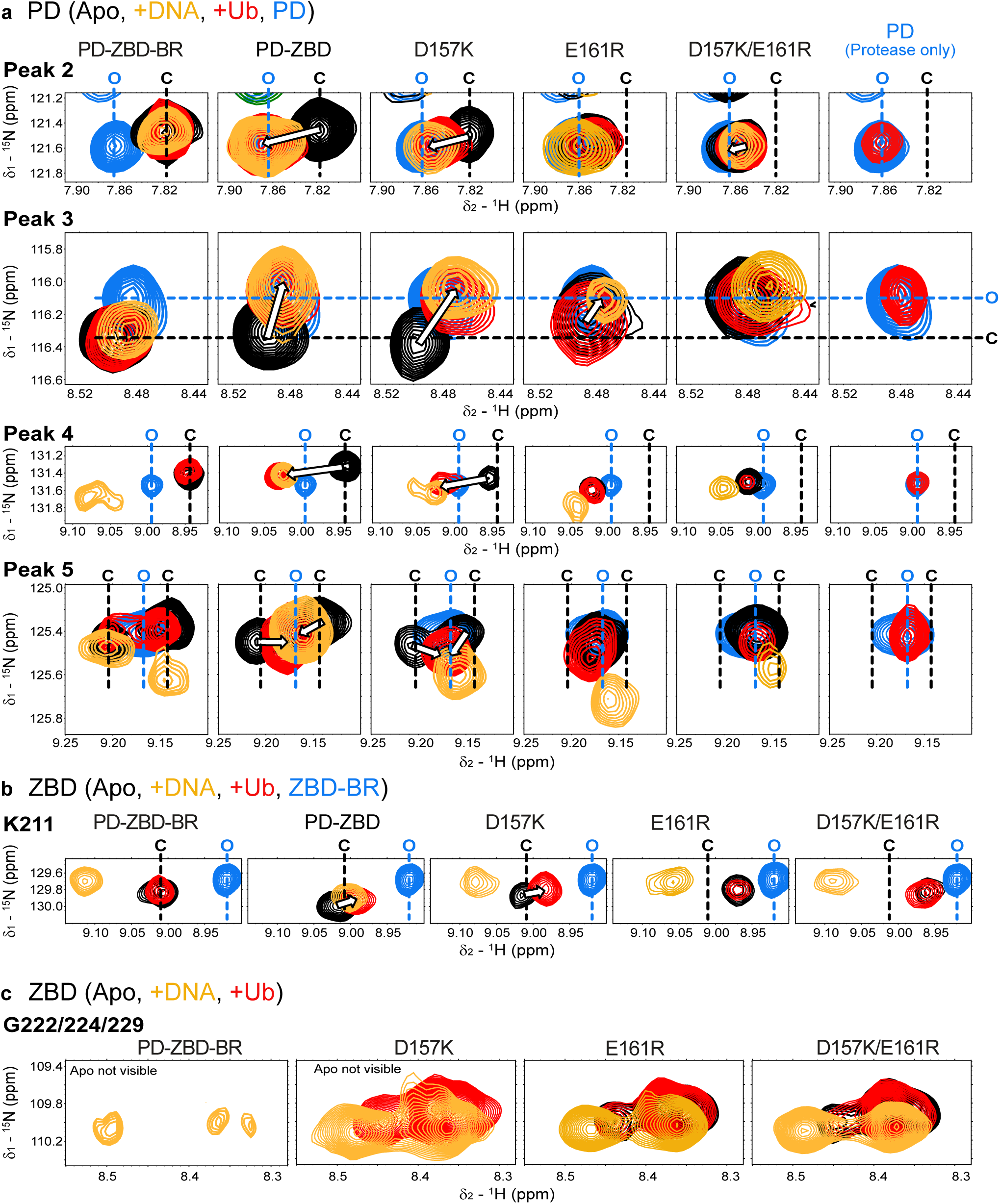
DNA- or ubiquitin-bindings shift conformations from closed to open states. a-c. Conformation-dependent resonances in the protease domain (PD) (**a**) (Peak 2-5, Supplementary Fig. 2), ZBD (K211) (**b**), and G222/224/229 (**c**), (see also Supplementary Fig. 2) of PD-ZBDD-BR (closed conformation), PD-ZBD, D157K, E161R and E157K/E161R, and PD (open conformation, for protease domain) or ZBD-BR (open conformation, for ZBD) comparing apo (black) and in the presence of a 5-fold molar excess of ubiquitin (Ub) (red) or a 2-fold molar excess of DNA (yellow). Open conformation resonances (PD for protease domain, ZBD-BR for ZBD) are shown for all in cyan.

